## Supplementary material for "ASPPs multimerize protein phosphatase 1": S3 File

**Supplementary File 1**

**Strains and Alleles**

**Figure 1A & 1B**

- GUN786	*mlt-4(mew193[mlt-4::tagRFP-T::HA]) V ape-1(mew342[APE-1::GFP::3xflag]) V*
- GUN1317 *mlt-4(mew193[mlt-4::tagRFP-T::HA]) V ape-1(mew422[APE-1(520-769)::GFP::3xflag]) V*
- GUN1551 *mlt-4(mew193[mlt-4::tagRFP-T::HA]) V ape-1(mew667[APE-1(133-248)::GFP::3xflag]) V*
- GUN1604 *mlt-4(mew193[mlt-4::tagRFP-T::HA]) V ape-1(mew719[APE-1(133-248,520-769)::GFP::3xFLAG]) V*

**Figure 2A**

- GUN1661 *mewSi262[Phyp7::TIR-1::GFP::3xflag] IV; mlt-4(mew118[MLT-4(E470K)]) V*
- GUN1662 *gsp-2(mew504[3xflag::AID::GSP-2]) III; mewSi262[Phyp7::TIR-1::GFP::3xflag] IV; mlt-4(mew118[MLT-4(E470K)]) V*
- GUN1663 *gsp-2(mew504[3xflag::AID::GSP-2]) III; mewSi262[Phyp7::TIR-1::GFP::3xflag] IV; mlt-4(mew193[mlt-4::tagRFP-T::HA]) V*

**Figure 3A**

- GUN666	*ape-1(mew342[APE-1::GFP::3xflag]) V*
- GUN810	*ape-1(mew378[APE-1(N583K)::GFP::3xflag]) V*
- GUN1064 *ape-1(mew379[APE-1(H591Y)::GFP::3xflag]) V*

**Figure 4B**

- N2 (Bristol)
- GUN350	*mlt-4(mew193[mlt-4::tagRFP-T::HA]) V*
- GUN713	*mlt-4(mew193[mlt-4::tagRFP-T::HA]) V ape-1(mew345[APE-1(N583K)]) V*
- GUN2049 *gsp-2(mew1004[6HNL(2mer)::GSP-2]) III; mlt-4(mew193[mlt-4::tagRFP-T::HA]) V ape-1(mew345[APE-1(N583K)]) V*
- GUN1954 *gsp-2(mew909[3HON(3mer)::GSP-2]) III; mlt-4(mew193[mlt-4::tagRFP-T::HA]) V ape-1(mew345[APE-1(N583K)]) V*
- GUN1956 *gsp-2(mew912[YRB1(4mer)::GSP-2]) III; mlt-4(mew193[mlt-4::tagRFP-T::HA]) V ape-1(mew345[APE-1(N583K)]) V*
- GUN1958 *gsp-2(mew914[6GWK(6mer)::GSP-2]) III; mlt-4(mew193[mlt-4::tagRFP-T::HA]) V ape-1(mew345[APE-1(N583K)]) V*
- GUN711	*mlt-4(mew193[mlt-4::tagRFP-T::HA]) V ape-1(mew346[APE-1(H591Y)]) V*
- GUN2094 *gsp-2(mew1004[6HNL(2mer)::GSP-2]) III; mlt-4(mew193[mlt-4::tagRFP-T::HA]) V ape-1(mew346[APE-1(H591Y)]) V*
- GUN1953 *gsp-2(mew909[3HON(3mer)::GSP-2]) III; mlt-4(mew193[mlt-4::tagRFP-T::HA]) V ape-1(mew346[APE-1(H591Y)]) V*
- GUN1955 *gsp-2(mew912[YRB1(4mer)::GSP-2]) III; mlt-4(mew193[mlt-4::tagRFP-T::HA]) V ape-1(mew346[APE-1(H591Y)]) V*
- GUN1957 *gsp-2(mew914[6GWK(6mer)::GSP-2]) III; mlt-4(mew193[mlt-4::tagRFP-T::HA]) V ape-1(mew346[APE-1(H591Y)]) V*

**Figure 4C**

- N2 (Bristol)
- GUN350	*mlt-4(mew193[mlt-4::tagRFP-T::HA]) V*
- GUN658	*mlt-4(mew193[mlt-4::tagRFP-T::HA]) V ape-1(mew340[deletion]) V*
- GUN2360 *gsp-2(mew1234[3HON(3mer)::GSP-2]) III; mlt-4(mew193[mlt-4::tagRFP-T::HA]) V ape-1(mew340[deletion]) V*
- GUN2364 *gsp-2(mew1238[6GWK(6mer)::GSP-2]) III; mlt-4(mew193[mlt-4::tagRFP-T::HA]) V ape-1(mew340[deletion]) V*

| **Allele** | **Generated by** | **Modified Locus** | **crRNA(s)** | **CRISPR repair** |
| --- | --- | --- | --- | --- |
| *mlt-4(mew118[MLT-4(E470K)]) V* | CRISPR (Beacham et al. 2022) |  |  |  |
| *mlt-4(mew193[mlt-4::tagRFP-T::HA]) V* | CRISPR (Beacham et al. 2022) |  |  |  |
| ape-1(mew340[deletion]) V | CRISPR (Beacham et al. 2022) |  |  |  |
| *ape-1(mew342[APE-1::GFP::3xflag]) V* | CRISPR (Beacham et al. 2022) |  |  |  |
| *ape-1(mew345[APE-1(N583K)]) V* | CRISPR (Beacham et al. 2022) |  |  |  |
| *ape-1(mew346[APE-1(H591Y)]) V* | CRISPR (Beacham et al. 2022) |  |  |  |
| *ape-1(mew378[APE-1(N583K)::GFP::3xflag]) V* | CRISPR | *ape-1* | rGB651 | oGB655 |
| *ape-1(mew379[APE-1(H591Y)::GFP::3xflag]) V* | CRISPR | *ape-1* | rGB656, rGB657 | oGB660 |
| *ape-1(mew422[APE-1(520-769)::GFP::3xflag]) V* | CRISPR (Beacham et al. 2022) |  |  |  |
| *gsp-2(mew504[3xflag::AID::GSP-2]) III* | CRISPR | *gsp-2* | rGB777 | 3xflag::AID for N-terminus of GSP-2 amplified with oGB778-oGB874 from pGB192 |
| *ape-1(mew667[APE-1(133-248)::GFP::3xflag]) V* | CRISPR | *ape-1* | rDTW282, rDTW401 | APE-1(133-248) amplified with oDTW499-oDTW500 from pDTW116 |
| *ape-1(mew719[APE-1(133-248,520-769)::GFP::3xFLAG]) V* | CRISPR | *ape-1* | rDTW581 | APE-1(133-248) amplified with oDTW582-oDTW583 from pDTW116 |
| *gsp-2(mew909[3HON(3mer)::GSP-2]) III* | CRISPR | *gsp-2* | rGB777 | 3HON amplified with oDTW801-oDTW802 from pYC26 |
| *gsp-2(mew912[YRB1(4mer)::GSP-2]) III* | CRISPR | *gsp-2* | rGB777 | Yrb1 amplified with oDTW805-oDTW806 from pYC29 |
| *gsp-2(mew914[6GWK(6mer)::GSP-2]) III* | CRISPR | *gsp-2* | rGB777 | 6GWK amplified with oDTW810-oDTW811 from pYC27 |
| *gsp-2(mew1004[6HNL(2mer)::GSP-2]) III* | CRISPR | *gsp-2* | rGB777 | 6HNL amplified with oDTW886-oDTW887 from pYC25 |
| *gsp-2(mew1234[3HON(3mer)::GSP-2]) III* | CRISPR | *gsp-2* | rGB777 | 3HON amplified with oDTW801-oDTW802 from pYC26 |
| *gsp-2(mew1238[6GWK(6mer)::GSP-2]) III* | CRISPR | *gsp-2* | rGB777 | 6GWK amplified with oDTW810-oDTW811 from pYC27 |
| *mewSi262[Phyp7::TIR-1::GFP::3xflag] IV* | CRISPR |  | rGB990 | Tir-1 amplified with oGB996-oGB997 from pGB210 |

**Plasmids**

| **Plasmid Name** | **Description** | **Generated by** | **Purpose** |
| --- | --- | --- | --- |
| pDTW123 | iASPP(602-828)::link(13)::HA::link(3)::TEV::Halotag in pcDNA5frt | Beacham et al. 2022 | Figure 2B |
| pDTW124 | iASPP(602-828, H665Y)::link(13)::HA::link(3)::TEV::Halotag in pcDNA5frt | Beacham et al. 2022 | Figure 2B |
| pDTW125 | iASPP(602-828, N657K)::link(13)::HA::link(3)::TEV::Halotag in pcDNA5frt | Beacham et al. 2022 | Figure 2B |
| pDTW131 | HA::link(3)::PPP1CA in pcDNA5frt | Gibson cloning | Figure 3B & 3C |
| pDTW231 | HA::link(3)::iASPP(602-828)::link(13)::TEV::link(5)::Halotag in pcDNA5frt | Gibson cloning | Figure 3B & 3C |
| pDTW232 | HA::link(3)::iASPP(602-828, N657K)::link(13)::TEV::link(5)::Halotag in pcDNA5frt | Gibson cloning | Figure 3B & 3C |
| pDTW233 | HA::link(3)::iASPP(602-828, H665Y)::link(13)::TEV::link(5)::Halotag in pcDNA5frt | Gibson cloning | Figure 3B & 3C |
| pDTW117 | empty::link(13)::HA::link(3)::TEV::Halotag in pcDNA5frt | Beacham et al. 2022 | Figure 3C |
| pDTW116 | APE-1(66-296)::link(13)::GFP::strepII in pUC57 | Gibson cloning | Template for CRISPR repair in mew667 |
| pYC25 | 3xflag::2mer(6HNL) | Chang and Dickinson 2022 | Template for CRISPR repair in mew1004 |
| pYC26 | 3xflag::3mer(3HON) | Chang and Dickinson 2022 | Template for CRISPR repair in mew909 and mew1234 |
| pYC27 | 3xflag::6mer(6GWK) | Chang and Dickinson 2022 | Template for CRISPR repair in mew914 and mew1238 |
| pYC29 | 3xflag::4mer(YrbI) | Chang and Dickinson 2022 | Template for CRISPR repair in mew912 |
| pGB192 | 3xflag::AID::link(3) in pUC57 | Gibson cloning | Template for CRISPR repair in mew540 |
| pGB210 | Pdpy-7::TIR-1::GFP in pUC57 | Gibson cloning | Template for CRISPR repair in mewSi262 |

**Oligonucleotides**

oGB655	GATGTCTCACAGGCCAATGATGAAGGGATCACGGCGTTGTACAATGCGATTTGTGCTGGA

oGB660	CCTCCATTTTCAGATCTCAGTTCTCAACGAAATGGCTCGATACATAAATCACATGTTCCG

oGB778	CTAATTGGGTTGTTTGAGCGATTTCTCGCAACAAATGAGTGACTACAAAGACCACGATGG

oGB874	GATATTGTCCAGATTAAGCTTTTCTACGTCTGTGCCACCCTTCACGAAC

oGB996	ATCAAAAATCCCAAAAACCAAAATGCAGAAAAGGATTGCATTGTCGTTTC

oGB997	GGAGACGCTTCCGCCGGTACCTCCACTGCCACCGCTTGTGCTGAGTCCGTTGGTGGTGAT

oDTW457	TCCCAGACTACGCCGGCGGAAGCTCCGACAGCGAGAAGC

oDTW458	GGCGTAGTCTGGGACGTCGTATGGGTACATGATGCTAGCCAGCTTG

oDTW499	TTCCCTCCATTTTCAGATCTCAGTTCTCAACGAGATGCCTTCTGAAATGATCGCCG

oDTW500	CGCTTCCGCCGGTACCTCCACTGCCACCGCTTGTGCTGGCTAGAAGAATCTGATGTTGC

oDTW582	CAGATCTCAGTTCTCAACGAGATGCCTTCTGAAATGATCGCCGATTAC

oDTW583	TCTTCTTTCAATCGCAACTTCCATTTCGAGGTTTTGGGCTAGAAGAATCTGATGTTGCTG

oDTW801	TCTAATTGGGTTGTTTGAGCGATTTCTCGCAACAAATGAGTGGATCCTCTGGAGTCCGTC

oDTW802	GTCCAGATTAAGCTTTTCTACGTCAGATCCTCCTCTTGGAAGTGGGGTGC

oDTW805	CTAATTGGGTTGTTTGAGCGATTTCTCGCAACAAATGATGGCTCAACAGAAGCTCG

oDTW806	TTGTCCAGATTAAGCTTTTCTACGTCAGATCCTCCCTGTCCCATAGACTTCACAGAC

oDW810	CTAATTGGGTTGTTTGAGCGATTTCTCGCAACAAATGATGTCTGCTGAGAAGAAGCA

oDTW811	TTGTCCAGATTAAGCTTTTCTACGTCAGATCCTCCATCGTCAGCATCAGCAGATG

oDTW886	AATTGGGTTGTTTGAGCGATTTCTCGCAACAAATGAGTGGCAGTCATATGGCTCATCAAC

oDTW887	ATTGTCCAGATTAAGCTTTTCTACGTCAGATCCTCCGAGTGAGGCCTTGACTCCA

oDTW1113	TCCCAGACTACGCCGGCGGAAGCCCCCAAAGTATGGAAATGCG

oDTW1114	GGCGTAGTCTGGGACGTCGTATGGGTACATGATGCTAGCCAGCTTG

**CRISPR guides**

rGB651	GGCCAATGATGAAGGGATTA

rGB656	GATCTCAGTTCTCAACGAGA

rGB657	CGAACAACAATGTCGAATAA

rGB777	GAGCGATTTCTCGCAACAAA

rGB990	ACCAAAATGAGCACAAGCGG

rDTW282	GATCTCAGTTCTCAACGAGA

rDTW401	AACAATGTCGAAAGTACTAG

rDTW581	CGAGATGCAGAATCTTGAAA
